# Key kinematic features in early training predict performance of adult female mice in a single pellet reaching and grasping task

**DOI:** 10.1101/2021.05.07.442851

**Authors:** Michael Mykins, Eric Espinoza-Wade, Xu An, Billy You Bun Lau, Keerthi Krishnan

## Abstract

Detailed analyses of overly trained animal models have been long employed to decipher foundational features of skilled motor tasks and their underlying neurobiology. However, initial trial-and-error features that ultimately give rise to skilled, stereotypic movements, and the underlying neurobiological basis of flexibility in learning, to stereotypic movement in adult animals are still unclear. Knowledge obtained from addressing these questions is crucial to improve quality of life in patients affected by movement disorders.

We sought to determine if known kinematic parameters of skilled movement in humans could predict learning of motor efficiency in mice during the single pellet reaching and grasping assay. Mice were food restricted to increase motivation to reach for a high reward food pellet. Their attempts to retrieve the pellet were recorded for 10 minutes a day for continuous 4 days. Individual successful and failed reaches for each mouse were manually tracked using Tracker Motion Analysis Software to extract time series data and kinematic features. We found the number of peaks and time to maximum velocity were strong predictors of individual variation in failure and success, respectively. Overall, our approach validates the use of select kinematic features to describe fine motor skill acquisition in mice and establishes peaks and time to maximum velocity as predictive measure of natural variation in motion efficiency in mice. This manually curated dataset, and kinematic parameters would be useful in comparing with pose estimation generated from deep learning approaches.

## Introduction

Early work in the 1980s established the use of the single pellet reaching assay for studying the acquisition of fine motor skill reaching in rodents (Evenden & Robbins, 1984; Whishaw et al., 1986). Rodents initiate reaching by lifting their forelimb from the ground, rotating their paw, advancing forward, clasping the pellet with their digits and returning the pellet back to their mouth (Nica et al., 2018; Whishaw & Pellis, 1990). Rodents are usually trained for 2-3 weeks and their motor performance is reported as percent success after training. Traditionally, these studies used detailed qualitative assessments of the reach-to-grasp task by decomposing it into ten phases and manually scoring each phase independently (Gharbawie et al., 2005; Whishaw, 2000). However, these scoring systems require extensive time for analysis, training of individual rodents, and inter-observer validation to reduce observer bias and drift. Most studies used overly trained rodents for circuit analysis and focus on end-point features (percent success, total attempts, and numbers of errors) (Schaar et al., 2010). Objective measurements that inform quality of motion are needed to improve our understanding of how a more efficient motion is learned with training. Natural variation in rodent motor performance is usually not reported, but there is considerable variability across individuals, ages, strains, and different litters (O’Bryant et al., 2011). Recent advances in deep learning and machine learning software are providing rich pose estimation for rodent reaching studies. With these advancements there is a push to focus more on individual variability vs stereotypies in behavior. We believe manual tracking of individual rodent reaching is needed to validate generated pose estimation and provide context in healthy and diseased models of motor disorders. We sought to determine if known kinematic parameters of motion efficiency in humans could predict natural variation in fine motor efficiency in mice during fine motor learning of the single pellet reaching assay.

## Methods

Adult female wild-type C57BL/6 mice (8-10 weeks old) were either individually housed or group housed with littermates on a 12hr light-dark cycle (lights on at 07:00 AM) and received food ad libitum until start of the procedure. At the start of the procedure, mice were placed on a calorie-restricted diet, in order to achieve a 10 percent reduction in their body weight. Water access was maintained 24/7 throughout this period. During dietary restriction, the weights of the mice were monitored each day. Once mice reached ~90 percent of their original body weight (after 9-12 days), they were introduced to the single pellet reaching assay chamber. Each day mice were placed in a Plexiglas chamber (lwh (cm): 20.3 × 8.3 × 21) containing a narrow-slit opening (width: 5 mm) towards an automatic pellet dispensing platform operated by an actuator (**Fig. 1**). An automatic pellet dispenser releases a pellet which is then brought up on a platform containing an infrared sensor. Any motion detected by movement to retrieve the pellet by the mouse will result in closure of the slit by a sensor activated door and lowering of the platform containing the pellet. After 10 seconds, the platform is raised and the door lifts to allow continued attempts to retrieve the pellet. Mice were placed in the apparatus and given 10 minutes to retrieve as many pellets as they can. During this time, mice were recorded using a Blackfly USB 3.0 camera (FLIR Systems Integrated; Wilsonville, Oregon). Videos were collected in SpinView (FLIR Integrated Systems) at 200 frames per second (FPS), with a 1000 μs exposure time and 628×647 pixel resolution. Observers recorded and scored each attempt to retrieve a pellet as a success (the mouse successfully grasps the pellet and brings it to the mouth) or failure (the mouse attempts to retrieve the pellet but does not bring it back to the mouth). Behaviors were performed between 9 AM-6 PM and this process was repeated for a total of 4 days of exposure for 5 animals. All procedures were in accordance with the National Institutes of Health’s Guide for the Care and Use of Laboratory Animals and approved by the University of Tennessee-Knoxville Institutional Animal Care and Use Committee.

**Figure 1:**
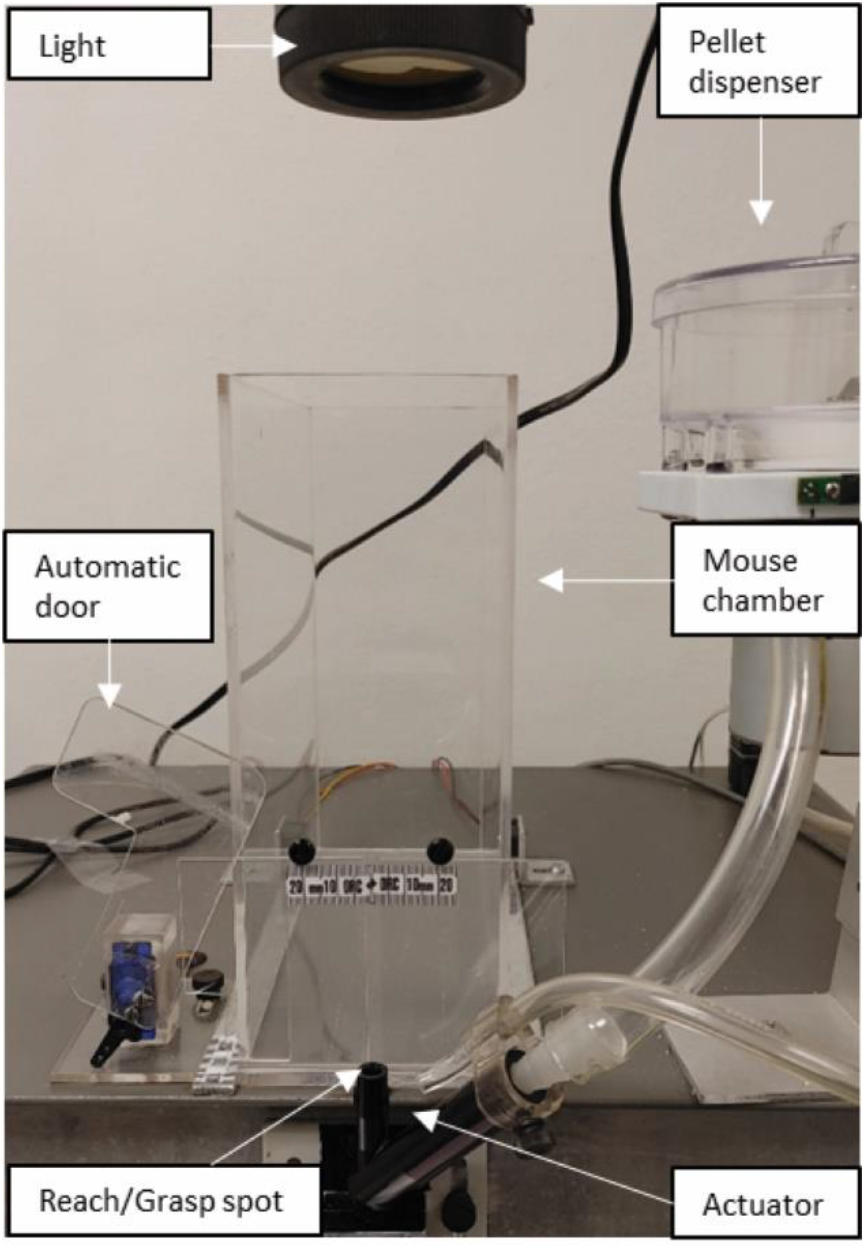
Representative picture of in-house motor apparatus. All major components of the motor apparatus are labeled within the diagram.

### Kinematic analyses of goal-directed reaching

Video data acquired during pellet reaching were imported into the free, open source Tracker motion analysis software (https://physlets.org/tracker/, Copyright © 2020 Douglas Brown, Robert Hanson, and Wolfgang Christian). All attempts, excluding errors (when a mouse reached in the absence of a pellet) were manually tracked and recorded from motion onset to offset. Motion onset was defined as the initiation of reach from a resting home position (i.e., the paw immediately before lifting from the ground). Motion offset was defined as the timepoint at which the pellet was brought to the mouth (successful attempt) or the return of the paw to the ground (fail attempt). Before analysis, a length scale was set in the Tracker by measuring the distance between the chamber opening and platform (~ 6 mm). Of the total 428 tracked reaches, 57 multiple reaches (3 or more attempts without returning the paw through the gate) were removed from further analysis as outliers. Video coders made note of the limb preference. Majority of mice used both limbs initially and emerged as left limb dominant. Paw location data (*x* and *y* position coordinates) extracted from Tracker were exported to MATLAB (R2019a) for further analyses. These extracted text files were named according to animal, type, trial number, and outcome (e.g., success or fail). Raw position data were differentiated to obtain velocity, acceleration, and jerk in the *x* and *y* directions. In addition, the resultant position, velocity, and acceleration were calculated using root mean square. A variety of kinematic features were then extracted from these time series data. The selected outcome measures included: the number of peaks, maximum velocity, time to maximum velocity, and mean squared jerk.

Measures were computed using standard definitions. The number of peaks were determined by summing instances when the derivative of the position data was equal to zero. Maximum velocity was determined by finding the maximum value from the resultant velocity time series. Time to maximum velocity was determined by subtracting the time of movement onset from the time corresponding to maximum limb velocity. Mean squared jerk was calculated using time normalized jerk:

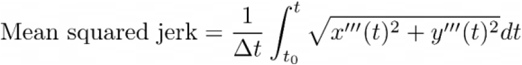

where *x*”’ and *y*”’ represent the third derivative with respect to time for the *x* and *y* paw coordinates (respectively), and *Δt* represents the time between sample frames. Analyses and data visualization were performed in R studio (R version 4.0.3). An unpaired Kruskal-Wallis with Dunn’s method was performed to test for statistical significance between number of peaks, time to maximum velocity, maximum velocity, and mean squared jerk of successful and fail reaches over days of exposure. Correlation plots, liner regression analyses, and sequence plots were generated in R studio to analyze the relationship between efficiency and days of exposure.

## Results

To determine if kinematic parameters used in human studies can predict motor learning efficiency in mice, we analyzed parameters during the initial exposure to the single pellet reaching task in adult female mice. These parameters include the number of peaks, maximum velocity, time to maximum velocity, and mean squared jerk. During the single pellet-reaching assay, mice were food restricted to increase motivation to reach to a high reward food pellet. Mice were recorded during behavioral performance, and pellet reaching was manually recorded using Tracker, an open source tracking software. We then derived relevant kinematic parameters to determine the relationship between these established parameters and motor efficiency in mice. We hypothesized that if patterns of motor output during a reach to grasp task were homologous between humans and rodents (Karl & Whishaw, 2013; Klein et al., 2012; Sacrey et al., 2009), then known human kinematic parameters can predict motor efficiency in mice.

The numbers of attempts, successes, and fails were manually scored to generate the average percent success efficiency across four days (**Fig. 2a**). On average, the percent success in adult female mice increased over time, which is consistent with previous observations in rats and mice (Buitrago et al., 2004; Chen et al., 2014). There was natural variability in performance during the early skill acquisition phase of single pellet retrieval. Some animals started out as fast learners or non-learners as previously defined (Chen et al., 2014). Animals 1 & 2 performed better initially and leveled off compared to animals 3 & 4, which started off poorly and got better with exposure. Animal 5 performed poorly and did not achieve a high percent success compared to the other animals.

**Figure 2:**
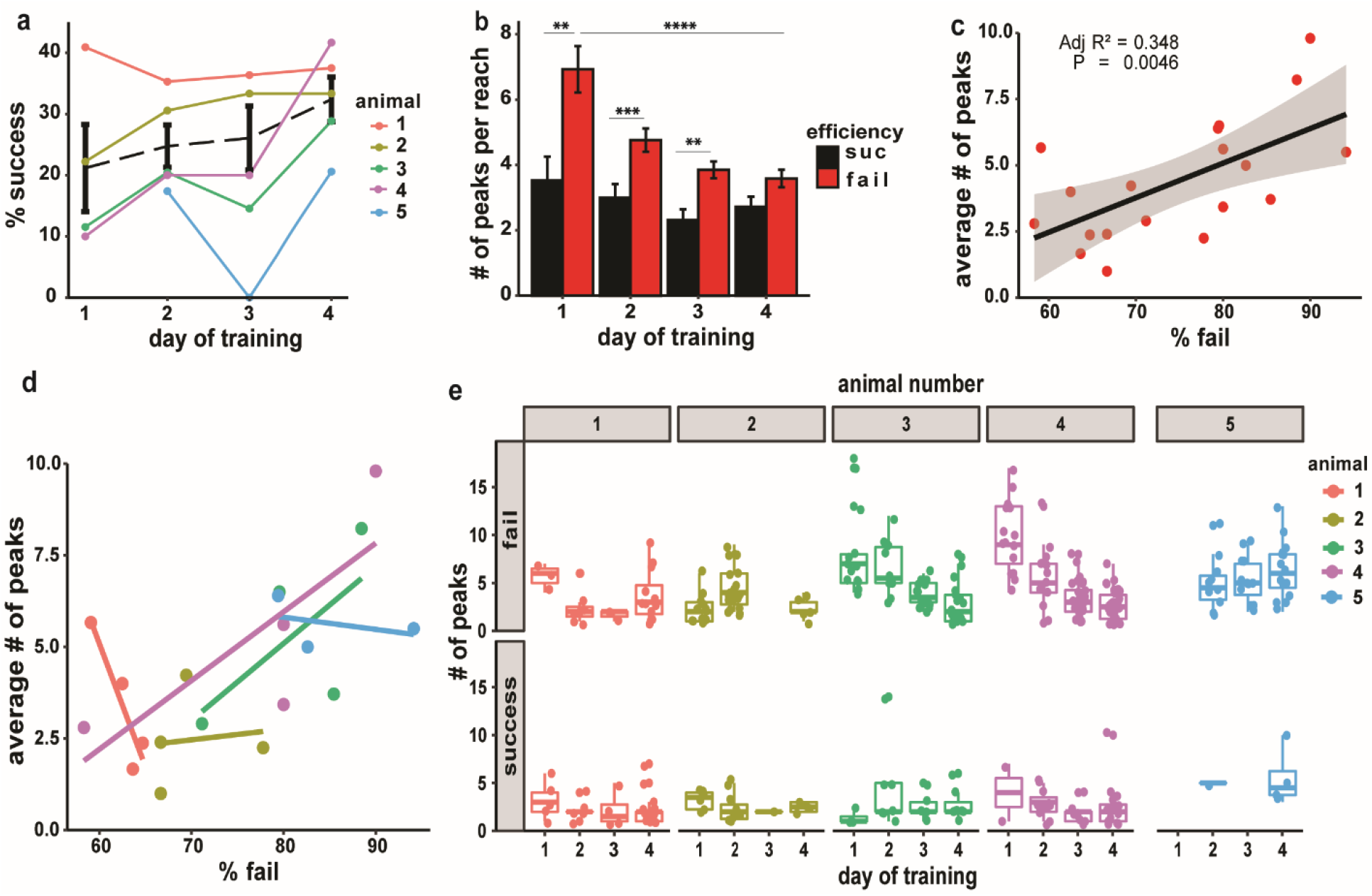
Peaks are a predictive measure for individual variation in percent failure measurement. **a.** Average percentage of successful pellet retrieval (number of success/total number of attempts) across training for n=5 mice (dashed black line). Mean ± S.E.M. of the percentage for each day of training. Animals are color coded by their percent success. **b.** Number of peaks per reach across training for both success and failed reaches during the pellet retrieval of 5 mice (n=378). Each bar with SEM represents the average number of peaks with standard error mean in reaching for that day. Statistical significance determined by non-parametric Kruskal Wallis followed by Dunn’s test, *P<0.05, **P<0.02, ***P<0.01, ****P<0.001. **c.** Linear regression analysis between the average number of peaks across training and percent failure. Each point represents the average number of peaks for failed reaches of one animal on a specific day. Line of best fit for proposed model and the 95 percent confidence interval is marked by the gray boundary; (r)= 0.5667, Variance (r^2^) = 0.3213, p-value = 0.0083. **d.** Individual correlation between average number of peaks and percent failure rate for each animal on a given day. **e.** Individual boxplots showing the distribution of peaks for each animal on each day across training. Box plots represent the median of each dataset.

In human studies, peaks (a measurement of movement intermittency) are predictive measures of motor efficiency (Balasubramanian et al., 2015; Berthier & Keen, 2006; Mancini et al., 2011; Rand et al., 2000; Teulings et al., 1997). Peaks in position, also referred to as sub-movements, are defined as in acceleration ‘zero-crossings’, between accelerating and decelerating phases of motion(Rohrer et al., 2002, 2004). The number of peaks is inversely related to efficiency and smoothness. In our study, the average number of peaks in successful reaching did not change with additional exposure (**Fig. 2b**). However, the average number of peaks in failed reaches decreased significantly between Day 1 (D1) and Day 4 (D4). By D4, there was not a difference in the average number of peaks between successful versus failed reaches, indicating a more efficient motion (**Fig. 2b**). Linear regression analyses showed a significant positive correlation between percent failure and the average number of peaks for failed reaches (r^2^=0.348, p-value < 0.01) (**Fig. 2c**). There was no correlation between percent success and the average number of peaks (r^2^=0.224, p-value > 0.05). Individual animal correlation of average peaks in failed reaches to percent failure was a strong predictor of natural variation in percent failure (**Fig. 2d**). To capture natural variation, we examined the distribution of peaks for failed reaches and successful reaches. The distribution of peaks for successful reaching did not change across days (**Fig. 2e, bottom panel**). The distribution of peaks in failed reaches decreased across days, most notably for animals 3 & 4 (**Fig. 2e, top panel**), also seen concomitantly with percent failure (**Fig. 2d**). For animals 1 & 2, we did not observe similar patterns, likely due to their relative efficiencies in successful reaching of the pellets, and relatively low number of peaks compared to other animals. The distribution of peaks increased with days for animal 5, consistent with inefficient pellet reaching. Thus, the more efficient a mouse is at pellet reaching, the more likely it is to have a low number of peaks in its reaching strategy. This result is consistent with observations in human studies that demonstrated peaks decrease with motor learning and improved motor function (Berthier & Keen, 2006)

In humans, failed attempts in a reaching task are followed by corrective strategies to increase success (Taylor et al., 2014). With continued attempts in trials, subsequent movements in a reaching task are less variable suggesting a corrective feedback mechanism for cognitively improving performance in reaching in humans and mice (Becker et al., 2020; Duthoo et al., 2014; Verstynen & Sabes, 2011). We plotted the sequences of number of peaks for all attempts during the 10-minute trial across days (**Fig. 3**). On D1, there is considerable variability in the sequence of number of peaks for all animals. On subsequent days, for animals 1-4, failed reaches with high number of peaks (warmer colors) are followed by attempts with a lower number of peaks (cooler colors). As we might expect, this observation was not observed in animal 5 which performs poorly at the task. Thus, mice may use sensory feedback from previous failed attempts to adapt a more efficient reaching strategy in subsequent trials.

**Figure 3:**
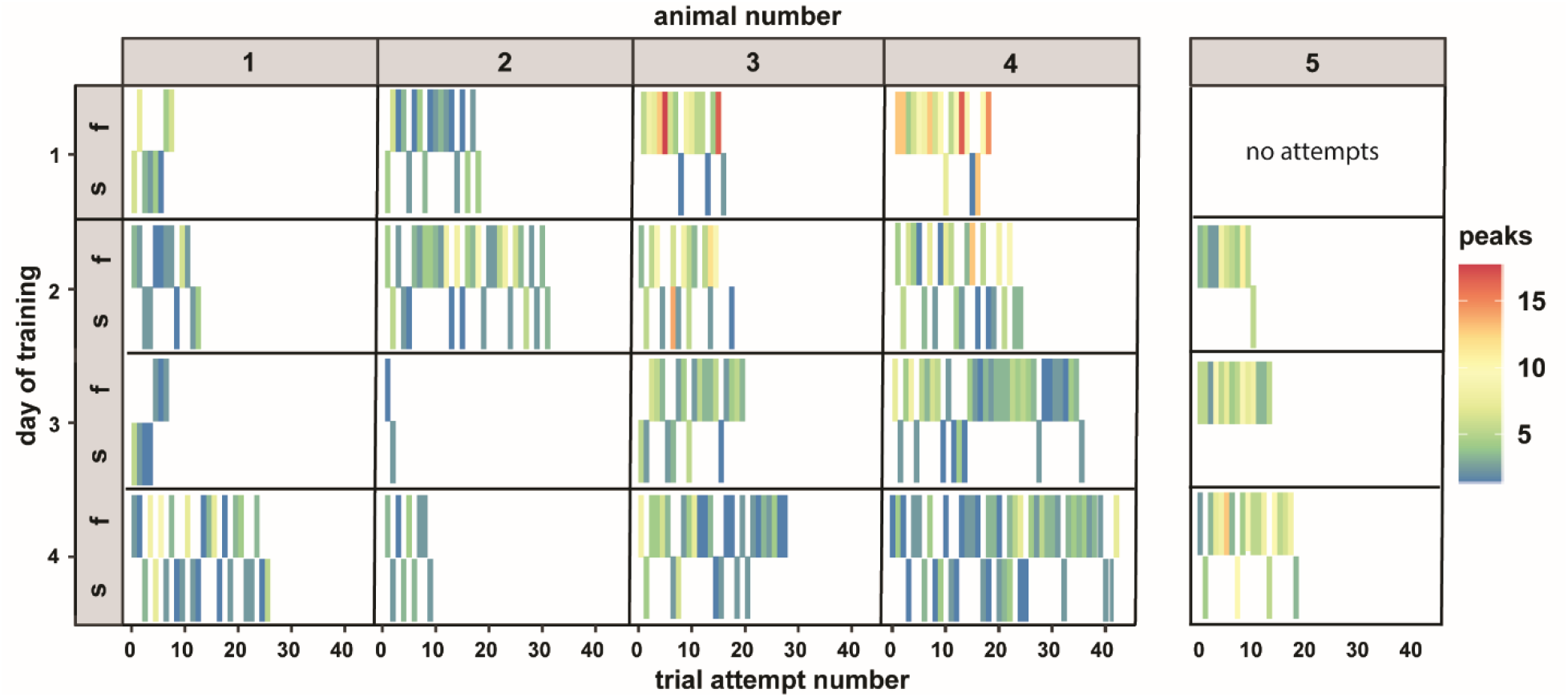
Mice correct the number of peaks in their reaching strategies after failure. Sequence analysis of number of peaks for all reaches during a trial on each day per animal. Number of peaks are denoted by a color scale; warmers colors indicate a larger number of peaks and cooler colors indicate a small number of peaks. Y axis represents the day of training grouped by efficiency and x axis represents the trial number of each reach grouped by animal.

In humans, time to maximum velocity and maximum velocity, also known as time to peak velocity and peak velocity, decrease as a motion becomes more efficient (Ma & Trombly, 2004; Wang et al., 2018; Wu et al., 2000). Maximum velocity of reaching did not change between success vs fail reaches in our mice (**Table 1.**). Time to maximum velocity (s) decreases significantly between D1-D4. Time to maximum velocity of successful reaches and fail reaches decrease significantly across days (**Fig. 4a**). Linear regression analyses showed a significant negative correlation (r^2^=0.3717, p-value < 0.01) between percent success and average time to maximum velocity (**Fig. 4b**) and a significant positive correlation between percent failure and average time to maximum velocity (r^2^= 0.4168, p-value <0.01) (**Fig. 4c**). These observations suggest that increased efficiency is correlated with a faster motion being adapted over time. Individual animal correlation showed average time to maximum velocity to percent success was a strong predictor of natural variation in percent success (**Fig. 4d**). To capture natural variation in motor performance, we analyzed the distribution of time to maximum velocity across each day for successful and failed reaches (**Fig. 4e**).

**Table 1.**
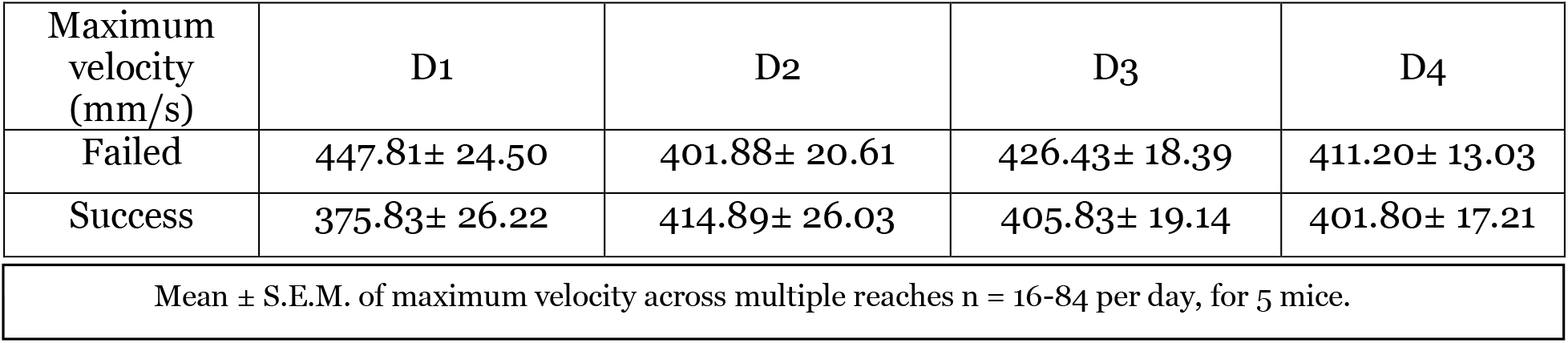
Average maximum velocity of failed and successful reaches across days.

**Figure 4:**
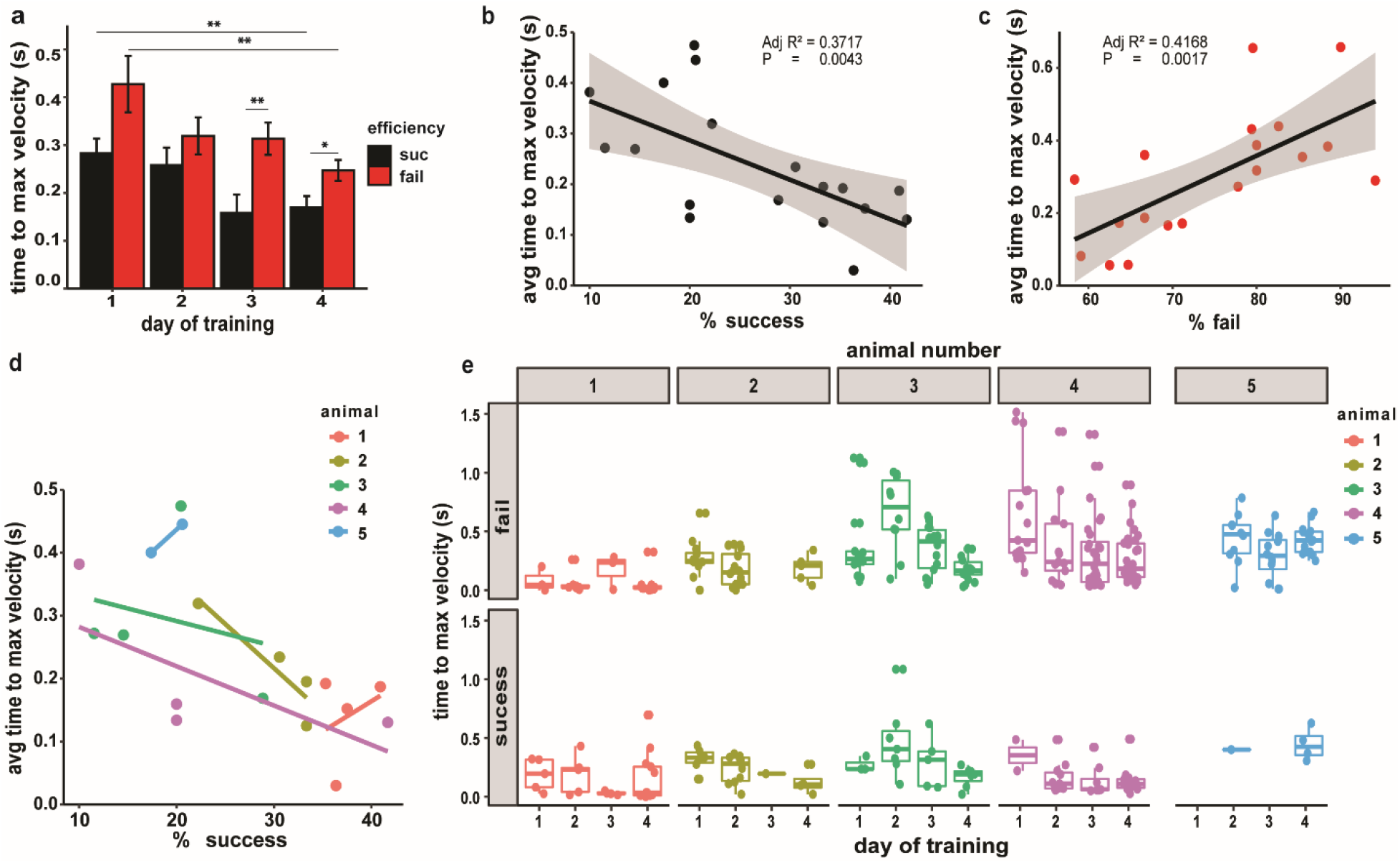
Time to maximum velocity is a predictive measure for individual variation in percent success a. Average time to maximum velocity across training for both success and failed reaches during the pellet retrieval of 5 mice (n=378). Each bar with SEM represents the average time to maximum velocity with standard error mean in reaching for that day. Statistical significance determined by non-parametric Kruskal-Wallis followed by Dunn’s test, *P<0.05, **P<0.02, ***P<0.01, ****P<0.001. **b, c.** Linear regression analysis between the average time to maximum velocity and percent success. Each dot represents the average time to maximum velocity of one animal on a specific day. Line of best fit for proposed model and the 95 percent confidence interval of the model is marked by the grey bands; for success (r)= 0.6159, Variance (r^2^) = 0.3739, p-value = 0.0042. **d.** Individual correlation between average time to maximum velocity, and percent success rate for each animal. **e.** Individual boxplots showing the distribution of time to maximum velocity for successful and failed reaches for each animal on each day. Box plots represent the median of each dataset.

Time to maximum velocity for both successful and failed reaches decreased across days most notably for animals 2, 3 & 4 (**Fig. 4e**), which was also observed concomitantly with increased percent success (**Fig. 4d**). For animal 1 we did not observe similar patterns, likely due to its relative efficiencies in successful reaching of the pellets, and relatively fast time to maximum velocity compared to other animals. For animal 5 the time to maximum velocity for both successful and failed reaches was large compared to other animals and did not change across days consistent with its inefficiency in pellet reaching. Next, we plotted the sequences of time to maximum velocity for all attempts during the 10-minute trial across days (**Fig. 5**).

**Fig 5.**
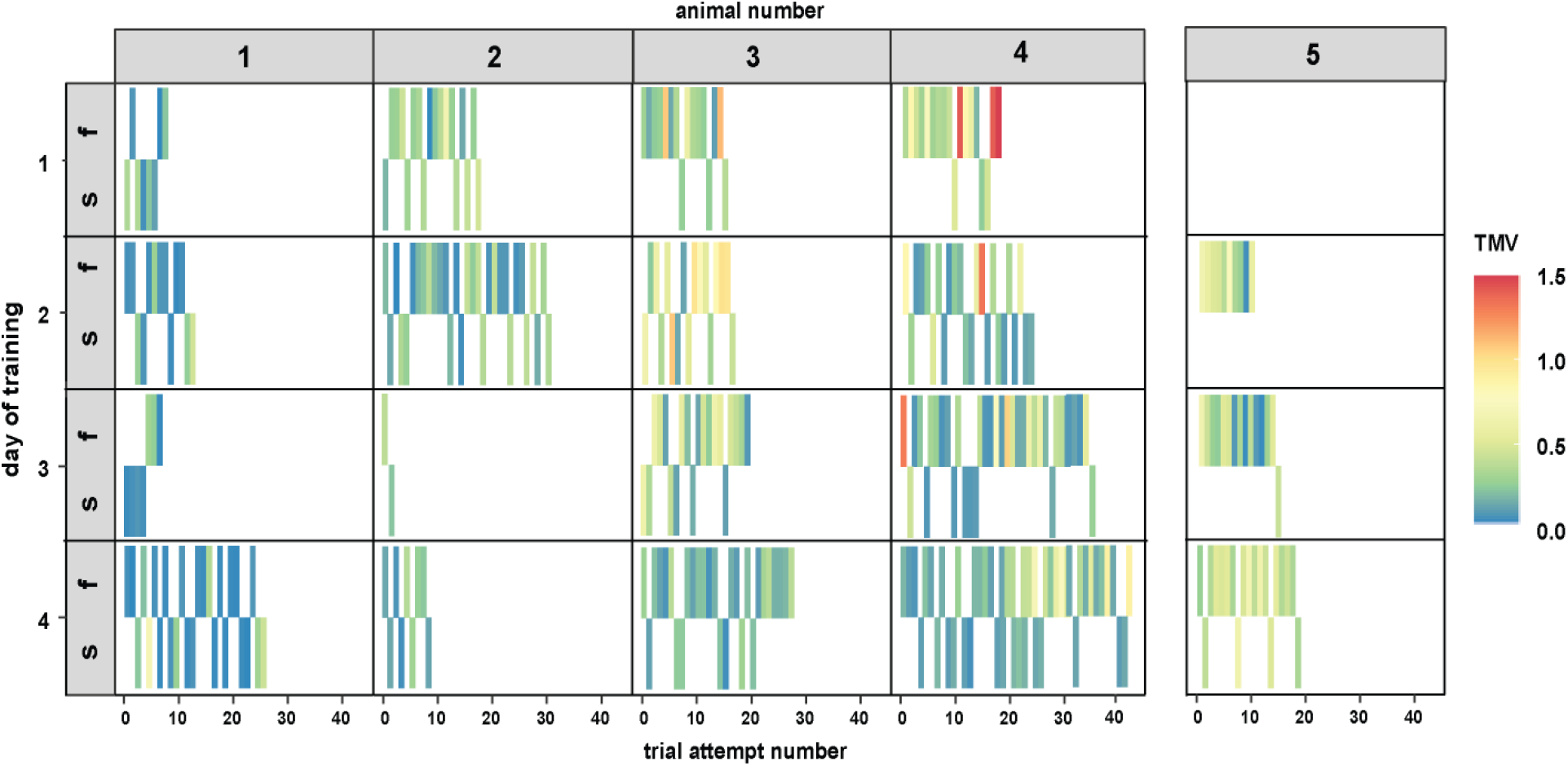
Mice do not correct the effort of reaching strategies after failure. Sequence analysis of time to maximum velocity for all reaches during a trial on each day per animal. Time to maximum velocity is color scaled (warmers colors indicate a slow time to maximum velocity and cooler colors indicate a fast time to maximum velocity). The y axis represents the day of training grouped by efficiency and x axis represents the trial number of each reach grouped by animal.

Sequence analysis of time to maximum velocity per reach between trials showed variability in the magnitude of time to maximum velocity for failed reaches (**Fig. 5, top panel**) and successful reaches (**Fig. 5, bottom panel**). Unlike peaks (**Fig. 3**), we did not observe corrective feedback between trials. Peaks in reaching decreased within trials, however, time to maximum velocity did not change within trials, although it was a strong predictor of success. We speculate this highlights a specific role for the timing of distinct neural circuits in shaping specific aspects of the motor output.

In humans, mean squared jerks of wrist and trunk motions increase in patients with Parkinson’s disease (Mancini et al., 2011; Teulings et al., 1997). Post-stroke individuals also exhibit increased mean squared jerk during reaching which is typically associated with a less efficient motion (Hogan & Sternad, 2009). In our study, there was significant increase in mean squared jerk (p <0.05) for successful reaches between D1 and D4 (**Table 2.**). There was a significant increase (p <0.05) in mean squared jerk for failed reaches between D2 and D4 (**Table 2.**). Linear regression analysis showed neither significant correlation between mean squared jerk and percent success nor percent failure.

**Table 2.** Average Mean Squared Jerk of failed and successful reaches across days.

Mean ± S.E.M. of mean squared jerk across multiple reaches $n = 16-84$ per day, for 5 mice. Bold values indicate significance with corresponding letter, a = \* $P < 0.05$ , b = \*\* $P < 0.01$ .
| Mean Squared Jerk ($\text{mm}^2\text{s}^6$) | D1 | D2 | D3 | D4 |
| --- | --- | --- | --- | --- |
| Failed | $3.60 \times 10^{13} \pm 2.48 \times 10^{12}$ | **$3.29 \times 10^{13} \pm 3.94 \times 10^{12,a}$** | $3.72 \times 10^{13} \pm 2.12 \times 10^{12}$ | **$4.43 \times 10^{13} \pm 2.85 \times 10^{12,a}$** |
| Success | **$3.39 \times 10^{13} \pm 3.94 \times 10^{12,b}$** | $4.19 \times 10^{13} \pm 3.12 \times 10^{12}$ | $4.36 \times 10^{13} \pm 3.35 \times 10^{12}$ | **$4.72 \times 10^{13} \pm 3.17 \times 10^{12,b}$** |

## Discussion

Since the introduction of the single pellet-reaching assay in 1980s, many groups have contributed to our knowledge on the neural circuitry required for fine motor skill execution. Previous studies validated the application of qualitative and quantitative kinematic parameters to assess recovery of motor function in stroke models of rats and mice, however, it is unclear if human kinematics are sensitive to acquisition of the skill (Berthier & Keen, 2006; Braun et al., 2012; Clarke et al., 2007; Lai et al., 2015). Clinicians assess motor efficiency by using scoring systems combined with kinematic parameters such as peaks, mean squared jerk, and time to maximum velocity (Bosecker et al., 2010; Carr et al., 1985; Fugl-Meyer et al., 1975). Scoring systems in rodents are not ideal as these systems rely on the presence or absence of motor deficits and cannot discern more subtle changes in movements in rodents (Sindhurakar et al., 2019). Growing evidence from qualitative and quantitative analyses of reach kinematics supports the notion that motions utilized by rodents and humans in reach-to-grasp task are homologous (Becker et al., 2020; Buitrago et al., 2004; Karl & Whishaw, 2013; Klein et al., 2012; Sacrey et al., 2009; Whishaw et al., 1992). Fine details of the reach-to-grasp motion in healthy mice are needed to identify subtle changes induced by motor perturbations, and to decipher motor learning sequences.

Our findings identify kinematic parameters that are sensitive to subtle changes in movements during fine motor skill acquisition in mice. According to our results, the numbers of peaks is a predictive measure of motion efficiency in mice. Peaks of failed reaches decreased with exposure and were comparable to peaks of successful reaches on D4. Peaks of failed reaches were a strong predictor of natural variation in percent success retrieval. We speculate peaks of successful reaches did not change significantly with training due to a maximum biological or mechanical efficiency. Time to maximum velocity decreased significantly with exposure and was a strong predictor of natural variation in success.

Furthermore, our results indicate a correlation between increased percent success and percent failure and efficiency of movements with short time to maximum velocity and minimal peaks, respectively. Sequence analysis suggests animals used performance feedback to alter reaching strategies between trials; these alterations were apparent in the number of peaks, but not time to maximum velocity, suggesting distinct neural circuits may shape the speed and efficiency of a motion at different time scales in response to corrective feedback after success and failure, respectively.

Dopaminergic circuits in the striatum of mice are important for modulating reach velocity, however it is unclear what circuits modulate sub movements in reaches (Panigrahi et al., 2015). The identity of these circuits may be related to specific neural circuits important for controlling the extrinsic (reaching) and intrinsic (grasping) properties for successful pellet retrieval (Azim et al., 2014; Esposito et al., 2014). One limitation from our study is our analysis focused on applying kinematics to the reaching component of this assay and did not focus on the grasping component. Failed trials could be due to animals not grasping properly in some trials, but these potential instances were not excluded from our analysis. As both the execution of reaching and grasping are important for successful retrieval, future work quantifying natural variation in mouse reaching and grasping kinematics will be required to investigate the underlying neural circuitry and its substrates.

We found that mean squared jerk was not a predictive measure of fine motor skill acquisition. Previous studies used mean squared jerk to quantify ballistic motions with set start positions. We reason that mean squared jerk is not an accurate measurement for this assay as free moving mice may start at any variable position and mean squared jerk is not sensitive to this variability (Becker et al., 2020). It is still possible that mean squared jerk may be a strong predictor for the acquisition of fine motor skills in head fixed mice, as the consistent start position mimics smooth target directed movements from previous human studies.

Assays such as rotarod performance test, forelimb hang, single pellet reaching assay are traditionally used to study fine motor learning. Licensed software is available to extract fine motor details from these behaviors; however, the amount of time and resources required for extracting fine details of motor behavior is substantially high. New approaches using deep learning allow for fast, automated, marker-less, pose estimation of free moving behavior in rodents (Graving et al., 2019; Mathis et al., 2018; Nath et al., 2019; Pereira et al., 2019). Mouse movement kinematics such as peaks, time to maximum velocity and more quantitative parameters can be quickly extracted from robust pose estimation data generated by DeepLabCut, DeepPoseKit, LEAP and SimBA. Our findings provide accessible quantitative measurements that can be used in future studies of fine motor skill acquisition and consolidation in healthy condition and models for neurological diseases.

In conclusion, through manual analysis of individual reaches during fine motor skill acquisition, we show that kinematic parameters identified from human studies such as peaks and time to maximum velocity are strong predictors of improved motor efficiency in adult mice. Our findings highlight kinematics that are sensitive to natural variability in fine motor skill acquisition which could reflect individual changes in the neural circuitry and neural substrates required for motor skill learning.

## Acknowledgements

We would like to thank the UTK Biology Service Facility (BSF), especially the late David Kidwell and Logan Sims, for engineering and maintaining the automatic pellet dispenser. We would also like to thank two undergraduate students, Andrew Cherosky and Carter Sanders for help in performing the experiments and gathering data. We would also like to thank Dr. Ralph Lydic for critical input on analysis. This work was supported by BCMB Departmental Graduate Fellowship awarded (MM), postdoctoral fellowship award from Rettsyndrome.org (BYBL), and startup funds from the University of Tennessee – Knoxville (KK).

